# Thermal Analysis of Protein Stability and Ligand Binding in Complex Media

**DOI:** 10.1101/2021.12.01.470796

**Authors:** Matthew W. Eskew, Albert S. Benight

## Abstract

Screening of ligands that can bind to biologic products of *in vitro* expression systems typically requires some purification of the expressed biologic target. Such purification is often laborious and time consuming and a limiting challenge. What is required, that could represent an enormous advantage, is the ability to screen expressed proteins in the crude lysate stage without purification. For that purpose, we explore here the utility of differential scanning calorimetry (DSC) measurements for detecting the presence of specific proteins and their interactions with ligands in the complex media where they were prepared, i.e. crude lysates. Model systems were designed to mimic analogous conditions comparable to those that might be encountered in actual *in vitro* expression systems. Results are reported for several examples where DSC measurements distinctly showed differences in the thermal denaturation behaviors of the crude lysate alone, proteins and proteins plus binding ligands added to the crude lysate. Results were obtained for Streptavidin/Biotin binding in *E. coli* lysate, and binding of Angiotensin Converting Enzyme 2 (ACE2) by captopril or lisinopril in the lysate supernatant derived from cultured Human Kidney cells (HEK293). ACE2 binding by the reactive binding domain (RBC) of SARS-CoV-2 was also examined. Binding of ACE2 by RBC and lisinopril were similar and consistent with the reported ACE2 inhibitory activity of lisinopril.

## INTRODUCTION

Production of biopharmaceuticals (biologics) and specific low abundance proteins using prokaryotic and eukaryotic expression systems provides an attractive alternative means for therapeutic development. Expression systems can provide for cheap and rapid production of a wide variety of biological compounds on various scales, including proteins, peptides, antibodies and small molecules [1]. These features enable production of otherwise low abundance proteins in sufficient quantities to serve as subjects for biochemical characterization and screen for ligand binding activities. For specific therapeutic applications, expression systems can provide for large scale production, on the metric-ton scale, of important drugs and vaccines [1]. Insulin is a classic example of a biologic produced on a large scale.

While expression of targeted biological compounds is fast and relatively inexpensive, execution of purification procedures for targeted compounds and their isolation can be both time consuming and costly. For moderately abundant and soluble proteins conventional purification strategies can typically require multiple steps and days to complete. Depending on the molecule, a multitude of purification steps may be necessary. Purification protocols requiring as many as 27 steps and four days have been reported [2]. Because of the time investment and associated costs of purification protocols, purified low abundance proteins can be expensive. The incumbent costs of *in vitro* expression products have curtailed their practical affordability.

Meeting demands of the biologics market and production of a range of desired target compounds requires generation of multiple expression vectors. These result in production of a library of candidate biologics that then must be screened. A process that is analogous to that of small molecule drug discovery where synthetically produced candidate compounds are screened and tested for their efficacy and bioavailability or ADMET properties. However, unlike small molecule drug discovery, biologics must first be expressed and purified before being tested. The added costs for preparation, testing, and screening of a therapeutic biologic can greatly increase research and development expenditures compared to the standard small molecule drug discovery pipeline [3]. These costs aside, biologics have shown great promise in treating a number of health conditions including autoimmune disorders and some cancers, that might otherwise be untreatable with conventional chemotherapy drugs [4]. Of the over 50,000 therapeutic compounds currently in the clinical trial pipeline, approximately 40% are biologics [3].

The uncertain and potentially unique nature of how biologics function in the body, present additional challenges limiting their development. Standard testing of biologic compounds in conventional animal models is at best marginally effective. Complications can arise because biologic compounds tend to target specific receptors or epitopes; and toxicological information does not necessarily translate between species. As a result only a few primate species have proven to be viable testing models [5]. Thus, it is essential that early in the testing process information regarding the pharmacokinetic/pharmaco-distribution (PK/PD) of produced compounds be obtained. Such information is crucial to guide toxicological studies, and properly assess human risk [5]. The considerable purification requirements can pose barriers to development since each lead compound must first be expressed and purified prior to testing of binding affinity for a targeted receptor.

Targeted biologic products of an expression system are expressed in culture media. In initial stages of the preparation process, cells harboring the over-expressed target are lysed releasing the target into a crude cell lysate. Unpurified crude lysates containing over-expressed proteins are produced at the beginning of the preparation process of a biologic. In the generic process, an expression system contains a constructed coding element (plasmid or other suitable vector) that codes for the desired target biologic. Cells containing the vector that produce the biologic target are grown in culture until cells can no longer grow or reproduce and expire. Harvested cells are centrifuged, resuspended in buffer and lysed. The genetic material (DNA/RNA) is then removed through extraction and centrifugation. At this stage, depending on the biologic, the desired, expressed unpurified protein can reside in either the lysed cellular media, or in the culture supernatant. Conventionally, it is at this stage that the laborious, multi-step purification process begins. Usually a process that must be completed before meaningful biochemical and biological testing of overexpressed compounds can be performed.

The testing scheme reported here is applicable at early stages of biologic production and circumvents the necessity for isolation and purification of the expressed protein target. This allows for testing and screening of ligand binding activity for the expressed target protein prior to purification. In the process, analysis of ligand binding to expressed proteins can be performed in unpurified cell lysates. As described subsequently the process utilizes the signal produced by differential scanning calorimetry (DSC) as a tool for detection of ligand binding to expressed proteins in the complex milieu of both prokaryotic (*E. coli*) and eukaryotic (Human Embryonic Kidney cells, HEK293) liquid culture expression systems.

## MATERIALS AND METHODS

### Chemicals and Reagents

Standard *E. coli* lysate was purchased from Bio-Rad (Hercules, CA) and received as a lyophilized powder. *E. coli* lysate samples were prepared by resuspending the appropriate mass of powder in standard PBS Buffer. Standard buffer for all experiments contained 150 mM NaCl, 10 mM potassium phosphate, 15 mM sodium citrate adjusted to pH = 7.4 with hydrochloric acid. Samples of the HEK293 cellular supernatant, Lot number: 03U27020D; unpurified angiotensin converting enzyme 2 (ACE2) expressed in human embryonic kidney cells (HEK293) received in the preparation media Lot number: 02U2802LL; isolated, purified ACE2 Lot number: 04U27020GC; and purified receptor binding domain (RBD) from SARS-CoV-2, Lot Number:05U22020TWB were purchased from RayBiotech (Peachtree Corners, GA). All HEK293 solutions had a volume of 200 µL. Captopril and Lisinopril were purchased from Sigma Aldrich (St. Louis, MO). Streptavidin was from Alfa Aesar (Haverhill, CA); biotin from VWR (Radnor, PA).

### Preparation of Samples for DSC Measurements

*E. coli* lysate was prepared by resuspending powdered lysate in standard PBS buffer to make a 2.7 mg/mL lysate solution. The HEK923 cell lysate was the supernatant after initial centrifugation of the crude cell lysate and received in liquid form at unknown concentration. Working solutions had a total volume of 500 µL containing 400 µL of HEK293 lysate and 100 µL standard buffer. Streptavidin, captopril, lisinopril, and biotin were prepared in standard buffer. All samples were incubated for 24 hours at 4 °C.

### DSC Measurements

DSC melting experiments were made using a CSC differential scanning microcalorimeter (now T.A. instruments, New Castle, DE). Samples were prepared by adding specific reaction components to pre-prepared media. For DSC melting experiments, the sample heating rate was approximately 1 °C/min while monitoring changes in the excess heat (microwatts) of the sample versus temperature [6–8]. In their primary form, melting curves or thermograms are displayed as plots of changes in microwatts measured for the sample, versus temperature. Since all experiments involved investigating interactions of ligands with proteins using DSC, we were interested in testing the sensitivity or our approach to detect proteins and protein/ligand interactions in the background solution comprised of the crude cell lysate. Consequently, all DSC measurements were made on samples in the lysate media, *E. coli* lysate or HEK923 supernatant, where samples were prepared. Thermograms were acquired for the complex media background, several proteins, and mixtures of these proteins with ligands, i.e. drugs or other another protein.

### Data Reduction and Analysis

Where the protein concentration was known, further refinement and normalization of the data provided the familiar DSC melting curve in the form of a plot of excess heat capacity, *ΔC*_*P*_, of the sample as a function of increasing temperature. Plots of *ΔC*_*P*_ versus temperature are commonly termed DSC thermograms. If the precise mass (concentration) of expressed protein was not known, thermograms were alternatively displayed in more primary form as plots of microwatts (*μW*) versus temperature. Baselines of the raw *ΔC*_*P*_ or *μW* versus temperature thermograms for *E. coli* were determined using a three point polynomial fit, over the temperature range of the transition, that produced a nearly straight line from 40 to 110 °C. Baselines of the raw *ΔC*_*P*_ or *μW* versus temperature thermograms for HEK293 samples were determined by connecting a straight line between data points at 40 and 100 °C. This baseline was then subtracted from the raw curves, producing baseline corrected thermograms used for further comparisons and analysis. For all experiments thermograms of the media alone served as the background that was subtracted from thermograms of samples containing drugs and/or proteins in the same media. These difference thermograms provided enhanced visualization of sample-specific changes, corresponding predominantly to the ligand/protein interactions of concern.

## RESULTS

### Overview

Investigations were aimed at defining thermograms of the media background, and determining whether thermogram measurements have sufficient sensitivity to detect the presence of protein alone in media, and effects of ligand binding on the protein against the background of complex media. Results are reported for several examples conducted under different conditions.

### Streptavidin-Biotin

Initial investigations explored streptavidin-biotin binding in reconstituted *E. coli* lysate. Binding behavior was examined under several conditions meant to mimic what might be encountered in an actual expression system. Figures 1–2 present results of this initial study. The thermogram of the *E. coli* lysate alone served as the background against which streptavidin-biotin binding was examined. Along with that of the media (alone) thermograms for lysate solutions containing streptavidin in different amounts, and streptavidin plus two molar equivalents of biotin in the background media solution are also shown in Fig 1. For each condition at least two independent experiments were conducted. Error bars shown in Figs 1–2 indicate the experimental reproducibility, which is quite good.

**Figure 1:**
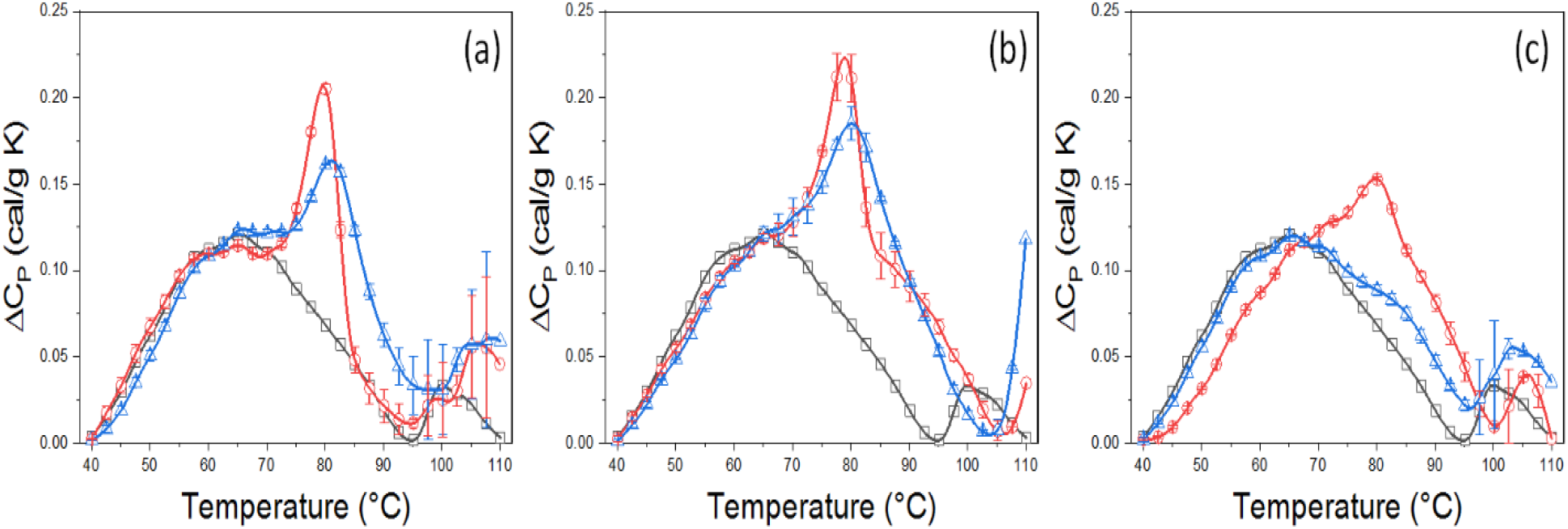
Thermograms of *E Coli* lysate spiked with streptavidin and streptavidin under different conditions. (a) Condition-I: 2 mg/mL *E Coli* lysate spiked with streptavidin to mimic 33% protein expression. (■) 2 mg/mL *E Coli* lysate standard. (●) 2 mg/mL *E Coli* lysate with 1 mg/mL streptavidin. (▲) 2 mg/mL *E Coli* lysate and 1 mg/mL streptavidin with 2 molar equivalents of biotin. (b) Condition-II: 1 mg/mL *E Coli* lysate spiked with streptavidin to mimic 33% protein expression. (■) 1 mg/mL *E Coli* lysate standard. (●) 1 mg/mL *E Coli* lysate with 0.5 mg/mL streptavidin. (▲) 1 mg/mL *E Coli* lysate and 0.5 mg/mL streptavidin with 2 molar equivalents of biotin. (c) Condition-III: 2 mg/mL *E Coli* lysate spiked with streptavidin to mimic ~11% expression. (■) 2 mg/mL *E Coli* lysate standard. (●) 2 mg/mL *E Coli* lysate with 0.25 mg/mL streptavidin. (▲) 2 mg/mL *E Coli* lysate and 0.25 mg/mL streptavidin with 2 molar equivalents of biotin. Error bars indicate experimental reproducibility for at least two experiments.

**Figure 2:**
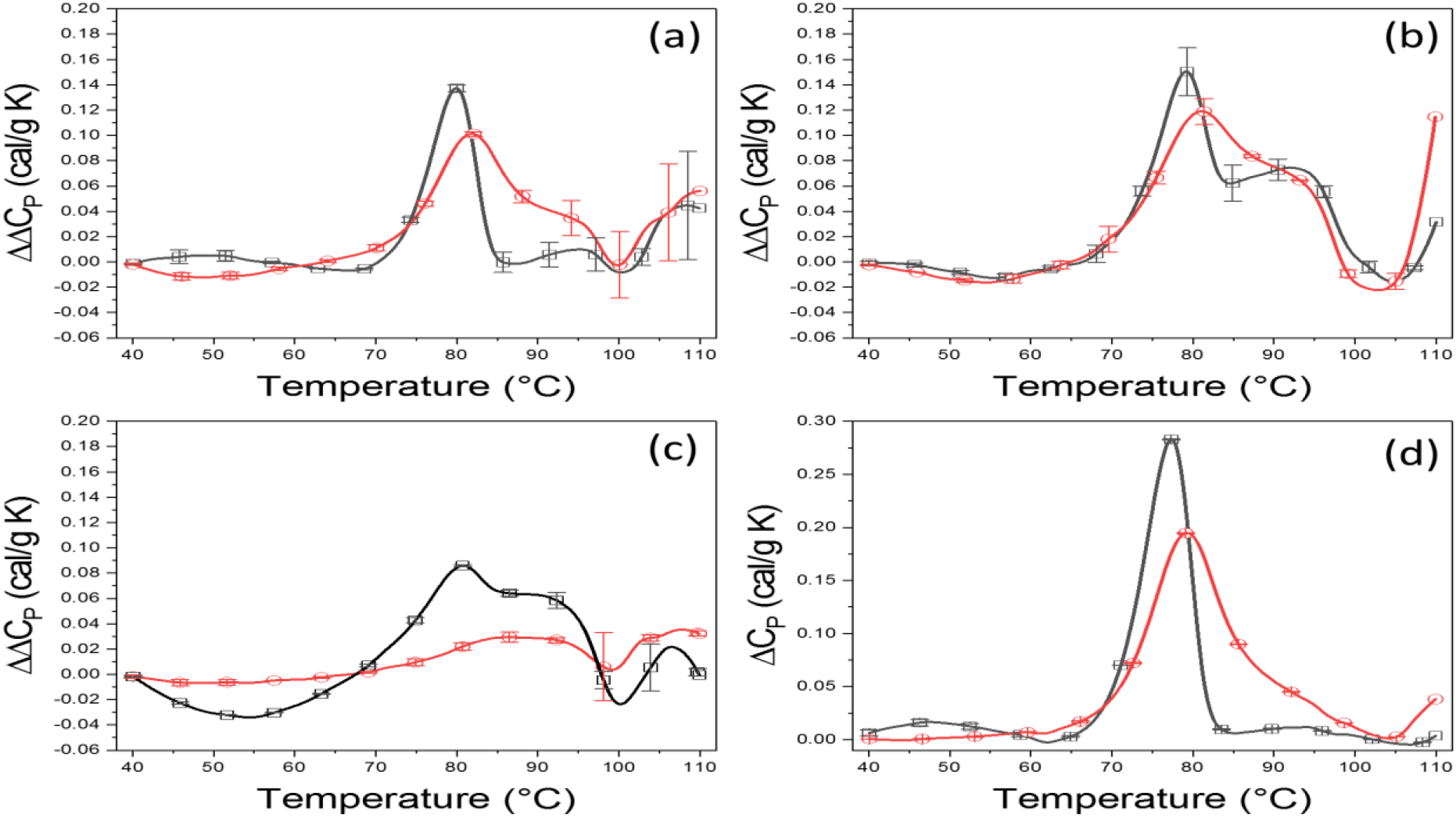
Difference Thermograms for streptavidin and streptavidin-biotin mixtures. (a) Difference thermograms for Condition I: The difference thermograms are comprised of the residual curve when the thermogram for *E. coli* lysate (2 mg/mL) alone was subtracted from the thermograms of *E. coli* lysate (2 mg/mL) + streptavidin (1mg/mL) (black line); and the thermogram of *E. coli* lysate (2 mg/mL) + streptavidin (1 mg/mL) + 2 molar equivalents biotin (red line). (b) Difference thermograms for Condition II: The difference thermograms are comprised of the residual curve when the thermogram for *E. coli* lysate (1 mg/ml) alone was subtracted from the thermograms of the *E. coli* lysate (1 mg/mL) + streptavidin (0.5 mg/mL) (black line); and the thermograms of *E. coli* lysate (1 mg/mL) + streptavidin (0.5 mg/mL) + 2 molar equivalents biotin (red line). (c} Difference thermograms for Condition III: The difference thermograms are comprised of the residual curve when the thermogram for *E. coli* lysate (2 mg/ml) alone was subtracted from the thermograms of the *E. coli* lysate (2 mg/mL) + streptavidin (0.25 mg/mL) (black line); and the thermograms of *E. coli* lysate (2 mg/mL) + streptavidin (0.25 mg/mL) + 2 molar equivalents biotin (red line). (d) Thermograms for Streptavidin (1 mg/mL) (black line) and Streptavidin (1 mg/mL) + 2 molar equivalents biotin (red line) in buffer. Note: shift of streptavidin thermogram (black line) with biotin binding (red line) similarly displayed a ~20% decrease in peak height as also seen in Figs 2a-2c.

Thermograms for streptavidin alone and the streptavidin/biotin mixture as shown in Fig 1 were measured under three different conditions. Condition-I was meant to mimic a 33% protein expression system. That is, the solutions contained 1 mg/mL of streptavidin added to 2 mg/mL of *E. coli* lysate. The streptavidin/biotin mixture contained streptavidin plus two molar equivalents of biotin in 2 mg/mL of *E. coli* lysate. Under Condition-I streptavidin concentration was half that of the lysate concentration making the total streptavidin concentration approximately 33% (w/w%). The three thermograms shown in Fig 1a correspond to Condition-I.

The 33% protein expression system was further investigated with half the concentrations of the components in Condition-I. For Condition-II samples contained 0.5 mg/mL of streptavidin added to 1 mg/mL of *E. coli* lysate; and 0.5 mg/mL streptavidin plus two molar equivalents of biotin in 1 mg/mL of *E. coli* lysate. Results for Condition-II are shown in Fig 1b. Even at half the concentrations of Condition-I, Condition-II displayed comparable signal resolution.

Examples thus far have demonstrated clear detection of ligand binding in a crude lysate at a 33% (w/w%) of total protein in the media (under Conditions -I and -II). While this value is not outside the range of possibilities, ~10% (w/w%) is likely a more realistic moderate level of protein expression [4]. To simulate this concentration range 0.25 mg/mL of streptavidin was added to 2 mg/mL lysate, corresponding to ~11% expression (Condition-III). Results are shown in Figure 1c. Comparison of the thermograms in Fig 1 clearly shows effects of biotin binding on the streptavidin thermogram, and sensitivity of the measurement to concentration. There are differences in Fig 1, but the trends of the thermograms and shifts with ligand binding are consistent under each of the three conditions. Under these conditions, different signals from streptavidin alone and streptavidin/biotin complexes in the lysate background are clearly detectable. While very good reproducibility was found between subsequent experiments over the range of 40 – 90 °C, results collected above ~100 °C displayed more variability probably due to slight aggregation in the lysate at high temperatures. Consistent with this supposition, variability over this range was more pronounced for samples containing 2 mg/mL lysate (Figs 1a and 1c) than for 1 mg/mL lysate (Fig 1b), as expected.

To enhance thermogram features of streptavidin and streptavidin/biotin complexes and their differences, difference thermograms where obtained by subtracting the thermogram for *E. coli* lysate alone (background) from the thermograms of streptavidin or streptavidin plus biotin. Results for Conditions I-III are shown in Fig 2. It would be expected that difference thermograms for Conditions-I and II, Figs 2a and 2b respectively, would be identical since thermograms are normalized by weight where *ΔC*_*P*_ is defined by cal/g K. However, there were slight differences between Figs 2a and 2b that warrant remark. The streptavidin peaks in Figs 2a and 2b display roughly equal peak heights at ~79 °C, also the addition of biotin causes a similar ~ 1 °C shift in Tm for the dominant peak, with a corresponding decrease in overall peak height. The primary difference occurs around the secondary peak that arises for streptavidin, in Fig 2b, at ~95 °C. This secondary peak was present and reproducible in all experiments under Condition-II. The source of this behavior is likely the lowered signal to noise ratio resulting from conversion of µW to *ΔC*_*P*_ under Condition-II; where small deviations above the background are magnified when accounting for the lesser mass of total protein in Fig 2b. That is, any deviations in Fig 2b would be magnified by 100% compared to those in Fig 2a because of the difference in total protein mass of the solution (3 mg/mL in Figs 2a and 2c versus 1.5 mg/mL in Fig 2b).

Difference plots determined at the relatively lower concentrations of Condition-III are displayed in Fig 2c. Clearly, the plots at 11% protein under Condition-III (Fig 2c) are somewhat different than those obtained at the higher 33% protein concentration (Conditions -I and -II, Figs 2a, 2b). Even so, in analogy to results for the 33% examples in Figs 2a and 2b, under Condition-III at 11% streptavidin, the temperature of the predominant peak (at ~80°C) is preserved but decreases in intensity at the lower concentration; while the peak at ~110°C increases. For reference, thermograms of streptavidin and streptavidin plus biotin alone in buffer (not lysate) are shown Fig 2d. These curves serve as the control for the isolated effect of streptavidin/biotin binding.

Fig 2d shows binding is typified by a temperature shift of 1.8°C, 20% reduction in peak height and increased width of the thermogram. Observations in Fig 2d are fully consistent with published calorimetric measurements of thermal denaturation of streptavidin [9]. Without biotin streptavidin has a melting temperature around 80 °C and shifts up from 1-2 °C under Conditions I-III.

Under all three conditions the shift of the thermogram of streptavidin in the presence of biotin is comparable. The 20% reduction in peak height for streptavidin-biotin alone, was consistent with those observed for streptavidin-biotin under Conditions I-III. The temperature shifts of streptavidin + biotin in buffer and under Conditions I-III are summarized in Table 1. The shift for streptavidin and streptavidin plus biotin in buffer is 1.8 °C. This shift decreases to 1.4 and 0.9 °C in *E. coli* lysate at decreased concentrations. Since this shift is the primary metric for ascertaining the presence of binding; the comparable temperature shifts summarized in Table 1 demonstrate the detection capabilities of the measurements, even at relatively lower concentrations.

**Table 1:** Temperature shift for Streptavidin + Biotin Thermograms for Conditions I-III

|  |  |  |  |
| --- | --- | --- | --- |
| Sample | Streptavidin $T_m$ (°C) | Streptavidin + Biotin $T_m$ (°C) | $\Delta T_m$ (°C) |
| 1mg/mL Streptavidin (control) | 77.39 ± 0.05 | 79.17 ± 0.06 | 1.79 ± 0.02 |
| 2mg/mL Streptavidin (33%, Condition-I) | 79.70 ± 0.02 | 81.16 ± 0.07 | 1.46 ± 0.05 |
| 1mg/mL Streptavidin (33%, Condition-II) | 79.03 ± 0.18 | 79.96 ± 0.36 | 0.93 ± 0.18 |
| 2mg/mL Streptavidin (11%, Condition-III) | 79.82 ± 0.10 | 80.71 ± 0.12 | 0.89 ± 0.02 |

The results with the streptavidin-biotin system clearly demonstrate sensitivity of the DSC measurements to detect presence of streptavidin and biotin binding in the *E*.*Coli* lysate; the directed primary purpose of the study. Differences between the thermograms for streptavidin alone, and streptavidin plus biotin in *E. coli* lysate were clearly discernable. Finding that binding of a ligand to a target protein in a crude lysate can be detected by DSC was a compelling preliminary result, and we were motivated by these results to venture further. However, the streptavidin-biotin system represents an ideal, and probably unrealistic, model of an expression system. Unrealistic because in the streptavidin-biotin system the total protein mass and mass percentage are precisely known prior to the experiment. Also, even in our moderate expression example (~11% w/w%) may still be higher than some or most real expression systems. In a real expression system, the actual amount of desired protein mass and percentage may not be known, or easily determined, particularly in early stages of development. In the next examples, products of a real expression system were used where concentrations of the expressed protein and lysate were unknown.

### Angiotensin Converting Enzyme (ACE2)

ACE2 expressed in liquid culture of human kidney cells (HEK293) was purchased from a commercial vendor. The supernatant containing ACE2 was provided from the vendor along with standard supernatant without expressed protein. The supernatant served as the background solution. Protein mass of ACE2 in the supernatant was unknown. Protein samples were provided in a specified volume, but the mass of ACE2 in that volume was also not known. Consequently, the actual mass of ACE2 in the sample solution, or what fraction of the total protein mass in the supernatant was comprised of ACE2, were unknown. This low level of information regarding the expressed protein in the supernatant likely resembles the situation that could be encountered in early-stage assessment of biologics expressed *in vitro*.

The thermogram measured for the supernatant, without ACE2 present, served as the background against which comparisons were made. Results are shown in Fig 3. Fig 3a displays the thermograms for the supernatant background and a solution of the supernatant containing an unknown amount of expressed ACE2. These curves have similar shapes but there is a clear difference for the thermogram of the supernatant containing ACE2. To isolate and enhance the specific contributions to the thermogram of ACE2 protein, the background thermogram for the supernatant alone was subtracted from that for ACE2 in the supernatant. The difference thermogram corresponding to ACE2 protein alone is displayed in Fig 3b; and shows two peaks at 55.6°C and 70.6°C. This observation is in precise agreement with published results of DSC measurements on unmodified somatic bovine ACE2, that also found two peaks at precisely the same temperatures. [10]. The peak at ~55°C was assigned to correspond to the C-domain of the protein; while the peak at ~70°C was assigned to the N-domain of ACE2 [10].

**Figure 3:**
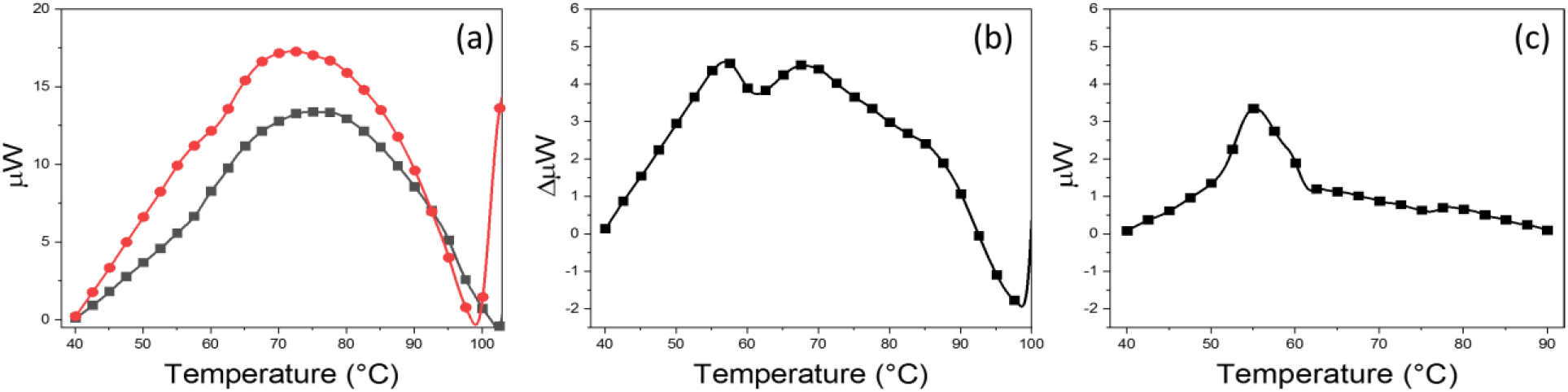
Thermograms for ACE2. (a) Thermograms for the HEK293 cell supernatant background (black line) and a solution of the supernatant containing an unknown amount of expressed ACE2 (red line). (b) The difference thermogram corresponding to ACE2 protein alone obtained by subtracting the curves in (a). (c) The measured thermogram for purified ACE2 in standard PBS buffer.

For comparison purposes, purified samples of isolated ACE2 were purchased and DSC thermograms were measured for those samples. The thermogram for the purified ACE2 that was purchased is shown in Fig 3c. When compared with the thermogram for unpurified ACE2 in Fig 3b, there are clear differences. Most notedly, the thermogram for purified ACE2 in Fig 3c does not match the thermograms for unpurified ACE2 in Fig 3b; nor does it match the DSC thermograms reported for somatic bovine ACE2 [10]. These differences between the measured thermograms for purified ACE2, and those measured for unpurified ACE2 and ACE2 reported in the literature [10] are remarkable.

To note, the purchased, unpurified and purified ACE2 purportedly contain only the extracellular domain. Interestingly in reference [10] modeling of ACE2 secondary structure from amino acid composition demonstrated that the N and C domains responsible for the dual thermogram peaks corresponded to the sequence from residues 1-734. In comparison, the expressed region for the purchased ACE2 was from Gln18-Ser740. Thus, this region should also have included both domains. It is possible that observed differences are due to the different origins of the ACE2 samples, bovine versus human. However remote, this possibility does not explain the presence of the extra peak at ~70°C in the expressed unpurified human ACE2.

These observations prompt several questions. If expression of Gln18-Ser740 is sufficient to suppress formation of the N domain peak at ~70°C, what are the possible origins of this peak on the thermogram of unpurified ACE2? The purification process could have affected the structure of ACE2, consistent with diminishment of the peak at ~70°C on the thermogram of the purified protein. That suggests the structure of the N-domain of the protein was somehow affected by purification. Also, if the serum-free cell supernatant of the unpurified ACE2 contains remnants of cellular ACE2 from the expression system, could these be responsible for the peak at ~70°C? This seems unlikely however since the sizes of the two peaks on the thermogram of unpurified ACE2 are essentially equivalent. If the cellular ACE2 exhibited normal dual peak behavior as reported in reference 10, in Fig 3b we would have expected to see a larger peak at ~55°C due to excess purified ACE2. But this was not observed.

Alternatively, the possibility exists that components in the unpurified sample bind to ACE2 thereby causing the observed dual peaks in the thermogram. We also view this as unlikely for the following reason. Previous results on ligand binding to proteins demonstrated that ligand binding is almost universally linked to changes of Tm on measured thermograms [7, 13]. Such Tm shifts associated with ligand binding are a well-known manifestation of Le Chatelier’s principle [13–15]. Since Tm for the two observed thermogram peaks are essentially identical to those reported in reference [10], ligand binding to ACE2 seems an unlikely explanation for the ~70°C peak. Additionally, ACE2 samples were purified using His-tag chromatography. If a ligand in the unpurified sample were to bind strongly enough to shift some portion of the ACE2 thermogram from ~55°C to ~70°C, we would have expected some ligand to remain bound and survive the chromatography purification step. As shown in Fig 3c for the purified sample, this is not the case.

Conventionally protein purity is usually confirmed using gel chromatography and measurements of protein mass by these methods are accurate so long as the mass does not change during the purification process. Such measurements might be expected to be largely insensitive to changes in structural integrity of the protein, possibly brought about by different steps in the purification process. Conversely, as evidenced by the thermogram in Fig 3, even though the mass of the protein was apparently unchanged, structural integrity might have been disrupted. Implications of this observation are highly significant for the following reason. Biologics, proteins or protein complexes, are intrinsically designed to target specific receptors. Much of the target specificity derives from critical interactions dictated by the overall structure in solution adopted by the biologic. If purification steps do significantly alter the biologic structure, activity of the compound could likewise be adversely affected, resulting in diminished efficacy.

For example, consider the following scenarios. Suppose the drug compound was designed to specifically bind the N-domain of ACE2, or expressed and purified ACE2 was used to test binding of drug compounds. If the N-domain of the ACE2 used in the binding assay was disrupted the binding activity could be considerably affected and provide a false negative result. Such a result in the context of screening could inadvertently lead to elimination of potentially active compounds. Additionally, if the N-domain were present but with a substantially altered structure (from truncated expression or purification), it is possible that the altered structure could actually have a higher binding affinity than the native N-domain structure. Again, leading to anomalous results, in this case a false positive. Although these are hypothetical scenarios, they highlight the unique utility of thermogram analysis and emphasize valuable insights in the screening processes, prior to protein purification. The aforementioned hypothetical scenario is especially relevant considering the numerous recently reported studies of SARS-CoV-2 (COVID-19); where the receptor binding domain (RBD) of COVID-19 specifically targeted ACE2 [16–20]. In which case damaged ACE2 used to assay binding, could provide spurious results.

### Binding of inhibitors to unpurified ACE2

Ligand binding measurements were also conducted on unpurified ACE2 in HEK293 cell supernatant plus added ligands. This next example involved a series of experiments aimed at detection of binding of known ligands to ACE2 in expressed, unpurified form, in the complex media background. Specifically, the ligands were captopril and lisinopril, both clinically active as ACE2 inhibitors to treat high blood pressure and heart failure. Thermograms were measured for solutions of ACE2 expressed in an unknown amount, plus captopril or lisinopril added to the supernatant.

Results are summarized in Fig 4. Thermograms of the lysate supernatant background, expressed ACE2 in the supernatant background, and the same solution plus captopril or lisinopril (at 250 µM) are shown in Fig 4a. In the media background all four curves have similar shapes, but also display clear differences. To better emphasize these differences, the thermogram of the background lysate was subtracted from the other three thermograms in Fig 4a rendering the difference curves shown in Fig 4b. These curves for ACE2 alone and ACE2+captropril and ACE2+lisinopril are clearly different, but there are some similar features. To further isolate and emphasize effects of the drugs alone on ACE2 stability, the background-subtracted thermogram for ACE2 alone in Fig 4b was subtracted from the background-subtracted thermogram of the mixture of ACE2+captropril and ACE2+ lisinopril (also Fig 4b). These difference curves are shown in Fig 4c. By removing contributions to the thermogram of both the supernatant background, and expressed ACE2, changes to ACE2 strictly due to effects of the ligands (captopril or lisinopril) on the ACE2+ligand thermogram emerge in Fig 4c. These isolated curves for lisinopril and captopril displayed in Fig 4c reveal remarkable differences between the two drugs. Lisinopril seems to have the largest effect on the region centered at 55 °C corresponding primarily to the C-domain region of the ACE2 thermogram; suggesting this may be the primary region of interaction between the drug and ACE2. As seen in Fig 4c, the difference thermogram of ACE2 containing captopril shows a relatively lesser effect on ACE2 compared to lisinopril. The difference thermogram for lisinopril suggests that, unlike captopril, lisinopril affects the stability of both the C and N domains of ACE2. These results for ACE2 and the ligands highlight different binding interactions for the two ligands that belong to the same class of drugs. Lisinopril is the stronger binder and has a greater effect on ACE2 activity than captopril. This observation is consistent with previous molecular docking studies that showed lisinopril is a stronger binder to ACE2 and has a higher affinity for the N-domain than captopril [18, 21, 22].

**Figure 4:**
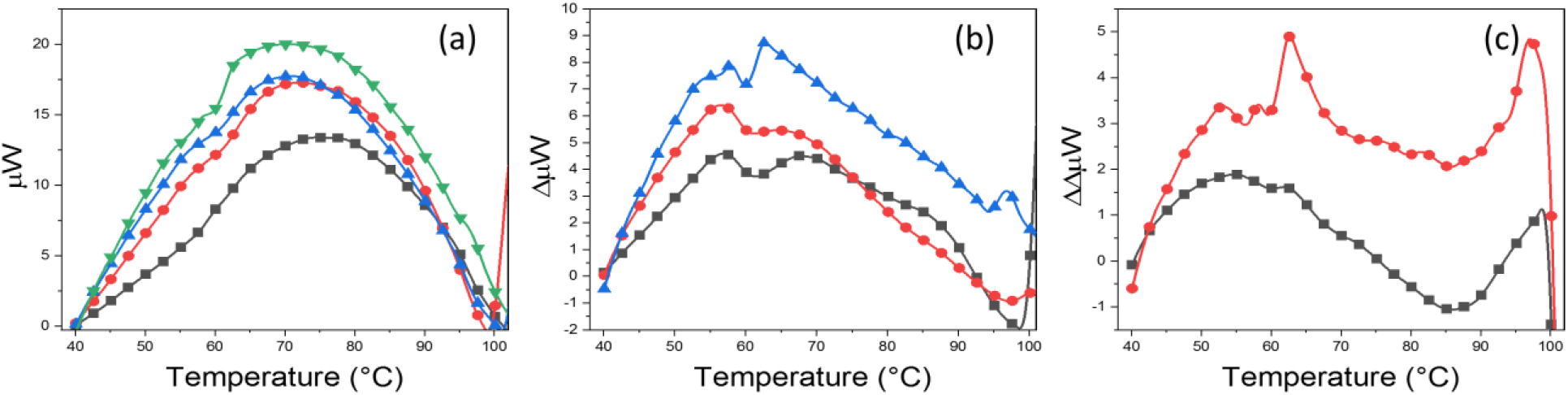
Thermograms of ACE2 with Ligands Lisinopril and Captopril. (a) Thermograms of the cell culture supernatant background (black line); expressed ACE2 in the supernatant (red line); expressed ACE2 in the supernatant plus captopril (blue line) or lisinopril (green line). (b) Differences of the thermograms in (a) after subtraction of the thermogram of the supernatant background. ACE2 (black line); ACE2+captropril (red line); ACE2+lisinopril (blue line). (c) Differences of the difference curves in (b) obtained after subtracting the curve for ACE2 alone from those for captopril (black line) and lisinopril (red line). These curves indicate changes to the ACE2 thermogram strictly due to effects of the ligand (captopril or lisinopril) interactions with ACE2.

### Interactions of COVID-19 RBD with ACE2

Examples described thus far involved analysis of systems meant to mimic small molecule drug ligand binding to expressed proteins. Experiments in the next example were designed to ascertain effects of protein-protein interactions with unpurified expressed protein in complex media. Such as may be the case when screening biologics which might be comprised of peptides or proteins that bind a particular expressed protein; or screening of potential protein targets for a specific protein ligand. A number of studies have determined the primary route of cellular infection by COVID-19 is binding of the viral RBD to ACE2 [16, 20, 23].

For this example, interactions of the RBD of COVID-19 with ACE2 were investigated. In this case, the RBD of COVID-19 was the ligand. Results are shown in Fig 5. Thermograms of purified RBD alone, unpurified ACE2 in the supernatant background and the purified RBD from COVID-19 added to unpurified ACE2 in the cell supernatant, are shown in Fig 5a. To better visualize contributions from RBD binding to unpurified ACE2 in the complex media supernatant, the thermograms of unpurified ACE2 in the supernatant and RBD alone in buffer were subtracted from the composite thermogram measured for the mixture of unpurified ACE2 and RBD. The resulting difference thermogram for the effects of the RBD protein ligand on unpurified ACE2 are displayed in Fig 5b (blue line). The largest intensity occurs between 50 and 60°C suggesting that that the interaction of RBD with ACE2 is primarily with the C-domain whose position shifts with binding.

**Figure 5:**
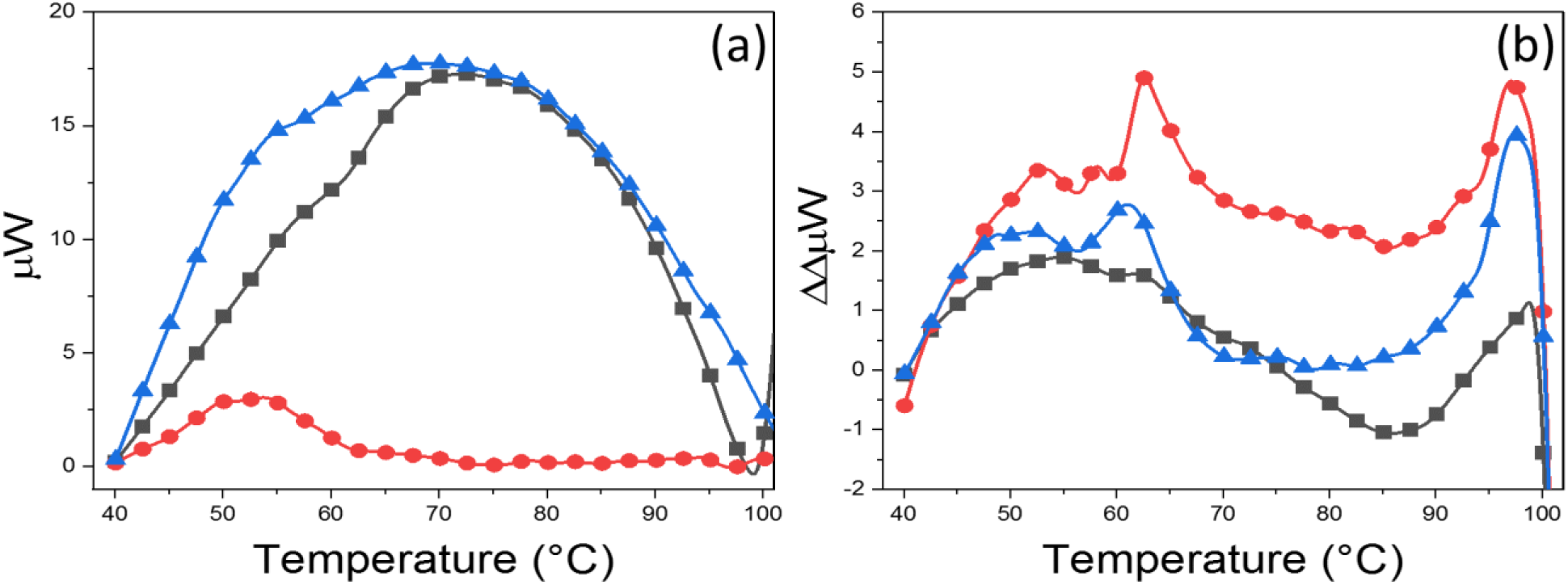
Thermograms of RBD from COVID-19. (a) Thermograms of purified RBD alone in buffer (red); unpurified ACE2 in the supernatant background (black); RBD added to added unpurified ACE2 in the supernatant background (blue). (b) Difference thermograms obtained by subtracting the thermograms of unpurified ACE2 in the supernatant and RBD alone in buffer from the composite thermogram measured for the mixtures of unpurified ACE2 plus RBD (blue line); ACE2 plus captopril (black line), ACE2 + lisinopril (red line). Note: the later difference curves for captopril and lisinopril were reproduced from Fig 4c.

### COVID-19, captopril and lisinopril interactions with ACE2

Since it has been determined that the route of COVID-19 viral infection is through binding of the RBD on the virus coat to the ACE2 protein, computational studies have been conducted for the purpose of designing or identifying drugs that can disrupt the ACE2/RBD interaction [19, 24–26]. In alternative approaches, researchers have also employed computational tools to virtually screen approved drugs for their potential binding to ACE2, and the RBD protein. Through this approach at least 40 commercially available compounds with suspected binding activity for ACE2 and RBD have been identified; offering them as possible inhibitors for the virus [19, 24]. However, experiments have been slow to confirm these computational predictions which currently amount to little more than theoretical conjecture. Confirmation has been slow mostly due to costs associated with testing compounds against the relatively expensive RBD and ACE2 protein samples.

The background subtracted thermogram for mixtures of ACE2 and RBD is shown in Fig 5b along with the difference curves for captopril and lisinopril. These results qualitatively demonstrate that, in terms of their thermogram responses, the interaction of COVID-19 RBD with ACE2 more strongly resembles lisinopril than captopril binding of ACE2. This observation might indicate that lisinopril, by binding strongly and in a similar manner to ACE2, may be an effective inhibitor of RBD binding. While lisinopril has proven ineffective at inhibiting COVID-19 infection, primarily by causing an upregulation of ACE2 in the body and subsequently providing more opportunities for infection; molecular modeling studies showed that based on binding characteristics lisinopril would be effective at inhibiting RBD binding to ACE2 [16, 19, 27].

## DISCUSSION

Results demonstrate the twofold application of DSC measurements for structural analysis of expressed proteins and rapid and sensitive detection of ligand binding of targeted proteins in complex culture media. This process provides a new means to probe the protein interactome and analyze complex solutions containing biologics produced by *in vitro* expression systems. For specific examples presented here attention focused on protein/small drug molecule interactions, but the process can be broadly expanded to interrogate protein-protein, antibody-antigen, protein-peptide, or interactions between any combination of biomolecules and small drug molecules.

Contrast this with conventional drug screening technologies (Isothermal Titration Calorimetry, Surface Plasmon Resonance, Mass Spectrometry, etc.) that require the use of highly pure compounds and solvents. Given the extreme cost and time associated with sample purification the ability to characterize “ impure” solutions and assess protein binding interactions provides a novel and powerful alternative for initial screening of new biologic compounds produced by *in vitro* expression. Results convincingly demonstrated detection of binding of small molecule drug compounds to proteins in unpurified lysate media. In a comparative manner, the process was able to detect ligand binding despite no precise knowledge of the protein composition of the HEK293 lysate e.g. total mass of protein in solution or the percentage of expressed protein. Applications for the capability of detecting binding in unpurified lysates are not limited to screening of biologic compounds. Based on results for detection of RBD binding from COVID-19 to ACE2 the analytical process provides a compelling application in the detection and analysis of viral outbreaks.

Several published studies reported DSC analysis of a number of different viruses showing diagnostically unique thermograms that provide an additional means of their characterization, [28–31]. Unique thermograms for different viruses could be due to differences in the array of proteins that typically decorate viral coat surfaces [30]. Such unique thermograms can also serve as the basis for screening biofluids such as plasma, mucus, cerebrospinal fluid, sputum, urine, etc) for the presence of live virus. Screening of the different biofluids combined with the analysis of expressed proteins in complex media enables a means for rapid screening of existing compounds for their efficacy in inhibiting binding to either protein targets on the virus, or the whole virus itself. Specifically, this has been an intense area of research activity during the COVID-19 pandemic where numerous computational studies have been undertaken in attempts to identify existing drugs that might disrupt the ACE2/COVID-19 RBD binding event by acting as a competitive inhibitor [16, 18, 19, 21]. Unfortunately, due to large resource requirements associated with acquiring purified ACE2 and COVID-19 RBD proteins, computational studies have not advanced past the *in silico* modeling (simulation) stage, or been experimentally verified.

Other than culture and amplification (both costly and time intensive) there currently are no methods available for accurate and rapid screening of biofluids for the presence of active virus. In this application, the DSC approach is positioned to perhaps make significant contributions. From our perspective, with the presence of virus particles in diseased biofluids comes the expectation that thermograms of infected samples will be significantly different compared to the normal thermogram for that biofluid. In this way thermograms can be extremely sensitive indicators of viral infection.

Previously the capture and retrieval of ligands from plasma, and subsequent confirmation of the identity of the retrieved analytes, was demonstrated for several different compounds [8]. While plasma is not the primary tissue compartment for COVID-19, it is the primary compartment for other infectious diseases such as human papillomavirus and hepatitis [32–35]. The capture process can be easily adapted to any biofluid or protein target. In principle, the actual physical isolation of virus particles directly from plasma, using the capture reagent, can be a far more accurate indicator of active infection than detection of remnants of viral RNA using PCR.

Although the above example focuses primarily on COVID-19, our approach is not limited solely to this virus. Being based on a physical measurement it provides a universal format and complimentary tool for analysis of virtually any disease (virus-based, or otherwise).

